# Atomic Resolution Mechanism of Ligand Binding to a Solvent Inaccessible Cavity in T4 Lysozyme

**DOI:** 10.1101/259549

**Authors:** Jagannath Mondal, Navjeet Ahalawat, Subhendu Pandit, Lewis E Kay, Pramodh Vallurupalli

**Affiliations:** Tata Institute of Fundamental Research, Hyderabad, India; Departments of Molecular Genetics, Biochemistry and Chemistry, University of Toronto, Toronto, Ontario, Canada; Hospital for Sick Children Program in Molecular Medicine, Toronto, Ontario, Canada

## Abstract

Ligand binding sites in proteins are often localized to deeply buried cavities, inaccessible to bulk solvent. Yet, in many cases binding of cognate ligands occurs rapidly. An intriguing system is presented by the L99A cavity mutant of T4 Lysozyme (L99A T4L) that rapidly binds benzene (~10^6^ M^−1^ s^−1^). Although the protein has long served as a model system for protein thermodynamics and crystal structures of both free and benzene-bound L99A T4L are available, the kinetic pathways by which benzene reaches its solvent-inaccessible binding cavity remain elusive. The current work, using extensive molecular dynamics simulation, achieves this by capturing the complete process of spontaneous recognition of benzene by L99A T4L at atomistic resolution. A series of multi-microsecond unbiased molecular dynamics simulation trajectories unequivocally reveal how benzene, starting in bulk solvent, diffuses to the protein and spontaneously reaches the solvent inaccessible cavity of L99A T4L. The simulated and high-resolution X-ray derived bound structures are in excellent agreement. A robust four-state Markov model, developed using cumulative 60 µs trajectories, identifies and quantifies *multiple* ligand binding pathways with low activation barriers. Interestingly, none of these identified binding pathways required large conformational changes for ligand access to the buried cavity. Rather, these involve transient but crucial opening of a channel to the cavity via subtle displacements in the positions of key helices (helix4/helix6, helix7/helix9) leading to rapid binding. Free energy simulations further elucidate that these channel-opening events would have been unfavorable in otherwise ligand-inactive wild type T4L. Taken together, by integrating experiments, these simulations provide unprecedented mechanistic insights into complete ligand recognition process in a buried cavity. By illustrating the power of subtle helix movements in opening up *multiple* pathways for ligand access, this work offers an alternate view of ligand recognition mechanism in a solvent-inaccessible cavity, contrary to common perception of *single* dominant pathway for ligand binding.

## Introduction

Measurements of affinities of small molecules to specific binding sites in proteins have become routine, especially in the context of drug discovery studies where such experiments are often among the first to be done. These thermodynamic measurements are routinely supplemented by kinetic studies of drug-receptor interactions [1–3]. However, in contrast to these standard kinetic and thermodynamic measurements, experiments that provide atomic level insights into the kinetic pathways by which small cognate molecules find their binding sites and the roles that protein conformational dynamics play in this process, are lacking. This is particularly the case for ligand binding sites that are deeply buried in the receptor protein core, precluding access, in the absence of structural rearrangements, even for bulk solvent molecules. Yet, many ligands recognize these deeply buried, solvent-inaccessible cavities very efficiently, and often at rates that approach those which are diffusion limited. Effective methodologies for the study of such binding processes in atomistic detail would have significant implications for the rational design of pharmaceuticals at a practical level and would provide important insights into molecular recognition and the role of protein dynamics in ligand binding, in general. representation, helix 4 (residues 83 to 91) in navy blue, helix 5 (92–113) in dark green, helix 6 (114–123) in red, helix 7 (125–134) in orchid, helix 8 (136–142) in light blue and helix 9 (142–156) in gold. B) The benzene molecule is not visible in the surface representation of T4L L99A because the binding site is buried in the protein with no pathway to the surface in the crystal structure.

The current work provides an atomistic view of the complete kinetic processes by which hydrophobic ligand binds rapidly to the occluded and solvent-inaccessible cavity of a well-known system namely the L99A mutant of lysozyme from the T4 bacteriophage (T4L L99A). Substitution of Ala for Leu at position 99 results in a 150 Å^3^ cavity [5] that can accommodate a range of ligands, including benzene, (Fig 1) [5–7]. Over the past several decades this protein has been used as a model system for understanding how buried cavities affect protein stability and structure, in general, and how ligands might navigate the protein landscape to bind rapidly to their proper sites. To this end, high-resolution X-ray structures of both the apo form of the protein and the benzene-bound complex have been solved by Mathews and coworkers [4] and pioneering studies have been undertaken that relate cavity size to protein stability [6]. NMR measurements have shown that the binding of benzene to T4L L99A is rapid, with an on rate of approximately 10^6^ M^−1^ s^−1^ [8]. Further NMR studies, based on relaxation dispersion approaches, established that T4L L99A exchanges in solution between a major conformer similar to the crystal structure of the protein and a minor state that is populated to 3% with a millisecond lifetime at room temperature [9,10]. However, the structure of this rare state shows that the cavity is occluded by the aromatic side-chain of Phe114 that prevents the binding of benzene so that the dynamic process that has been characterized is not relevant to ligand binding [11,12]. Oxygen and xenon binding studies have also been performed that have helped to identify hydrophobic cavities in the protein [13–15]. The system has also served as benchmark system for many binding free energy calculations.[16,17]

**Fig 1:**
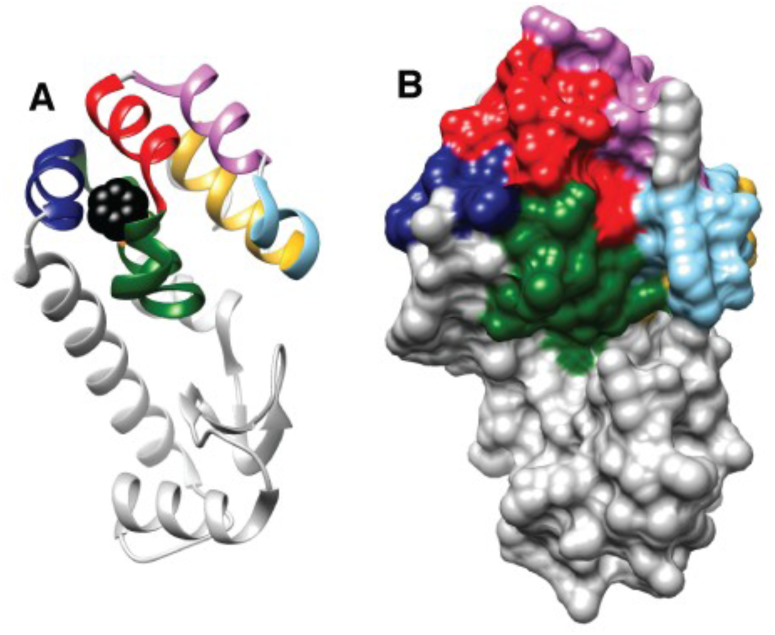
Benzene binds to a buried hydrophobic cavity in T4L L99A that is inaccessible from the surface. A) Ribbon representation of T4L L99A bound to benzene (PDB 3DMX) [4]. Bound benzene is shown in black CPK

Despite the availability of macroscopic kinetic rate constants for aromatic ligand binding, high resolution structures of the bound and free forms of T4L L99A, and NMR spin relaxation data that indicate conformational flexibility in the region surrounding the ligand binding site, a detailed description of how ligands bind to the protein remains elusive. Recent innovations in experimental and computational techniques, notwithstanding, the mechanism by which ligands bind to buried sites in a protein has remained difficult to address, because the process often involves the formation of transient metastable states that are challenging to characterize at high resolution and because the conformational fluctuations leading to binding can be rapid, complicating detailed characterization by most biophysical techniques. To this end, molecular dynamics simulations are emerging as a complementary tool to experiment for the study of molecular recognition processes because of recent innovations in GPU-based technologies, access to distributed computing, and the development of special-purpose computers [18–21], that have facilitated studies of many pertinent biochemical processes at atomic resolution.

In this work, we elucidate the mechanism by which benzene binds to the solvent-inaccessible cavity of T4L L99A by capturing the entire binding process using unbiased molecular dynamics (MD) simulations. Despite the fact that the simulations contained no *a priori* knowledge of the T4L L99A binding site, the resulting bound conformations that were obtained in a series of microsecond-long simulations and independent trajectories matched that of the crystal structure. Underlying Markov state model analyses [22–24] of our simulation trajectories established multiple binding pathways, each with transient opening of a channel involving a distinct pair of helices in the C-terminal domain of T4L L99A. Using an enhanced sampling technique, infrequently biased metadynamics simulations [25,26], the ligand unbinding pathways from the bound state have also been established. The resulting estimates for ligand binding on- and off-rates as well as binding free energies are in reasonable agreement with the experimentally reported values, lending credence to the conclusions of this study. Our work establishes that the fast binding of benzene to T4L L99A derives from its access to multiple pathways that are formed from low barrier concerted protein fluctuations that only occur in the mutated protein. It also establishes the utility of molecular dynamics simulations, in general, in providing detailed descriptions of binding processes that are subsequently validated by comparison of the resulting calculated kinetic and thermodynamic parameters with those obtained via experiment.

## Results

### Long unbiased simulations place benzene at the target-binding site of L99A T4L with atomic precision

We performed six independent all-atom unbiased MD simulations with lengths varying between 2 and 8 µs, resulting in a total simulation time of 29 µs. In each of the simulations a benzene molecule, starting from a random position in solution, was correctly placed in the target-binding site, corresponding to the hydrophobic and solvent-inaccessible cavity created by the L99A mutation in T4L. Ligands were initially positioned at least 4 nm away from the binding pocket in random orientations. As shown in three representative trajectories (Fig 2 and movies S1-S3 in Supporting Information), the ligand diffused extensively in the solvent, occasionally contacting different parts of the protein surface, before entering the binding pocket. Fig 2A quantifies the time-evolution of pocket-ligand distances en route to binding in three representative binding trajectories, where the distance between the binding cavity and benzene gradually decreases as the ligand finds its target. As depicted in Fig 2B, the MD derived structure converges to within 1–2 Å of the X-ray based model for the holo-form of the protein [4], as quantified by the root-mean-squared deviation (RMSD) of heavy atoms from residues defining the cavity and including the ligand (detailed in Materials and Methods). The excellent agreement between the simulated and crystallographic binding regions, including the orientation of benzene (Fig. 2C), and provides confidence in the underlying force field used to model the T4L L99A benzene binding process. Unlike standard docking approaches which search for the best ligand orientation within a predefined binding site, our long and unguided atomistic MD simulations reproducibly identified and maintained the correct ligand-bound pose without user intervention or incorporation of any prior knowledge of the binding site. As shown in Fig S1, in each of the trajectories substantial dehydration accompanies benzene entry into the hydrophobic and solvent-inaccessible binding pocket.

**Fig 2:**
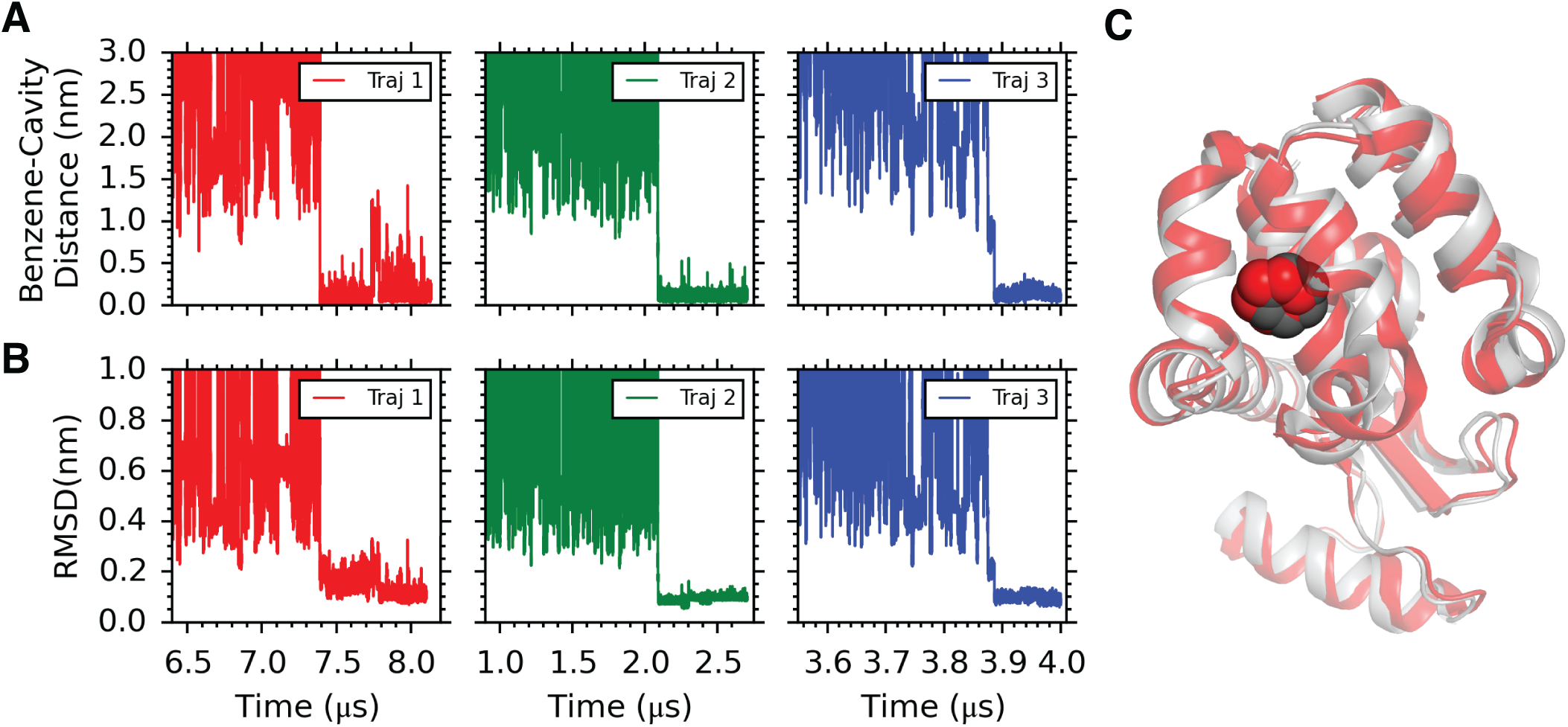
Benzene binds correctly to the L99A cavity in long unbiased MD simulations. A) Distance between the solvent-inaccessible cavity of T4L L99A and benzene shows that benzene diffuses through the solvent before finally binding. Protein heavy atoms within 5 Å of benzene in the holo-protein (PDB id: 3DMX) [4] are defined as the ‘binding pocket’ or ‘cavity’. Radial distances were computed between the respective centers of mass of the pocket and the ligand. B) Time profile of the root-mean-squared deviation (RMSD) between the MD derived structure and the crystal structure of the benzene-bound form of the protein. Heavy atoms from cavity residues plus ligand of the MD derived structure are compared with corresponding positions in the crystal structure of the benzene-bound form of the protein (PDB id: 3DMX [4]) to calculate the RMSD values. Initial RMSDs are high because benzene is not in the binding site and not because the protein is distorted. C) Overlay of a snapshot of the benzene-T4L L99A complex from simulations with the crystal structure (PDB id: 3DMX). The crystal structure is shown in grey with the benzene in black CPK representation while the protein in the MD snapshot is shown in red with the benzene in red CPK representation.

MD trajectories were further analyzed using Markov State Models (MSMs) (detailed later) to estimate kinetic and thermodynamic parameters and to obtain insights into the conformational dynamics that are responsible for binding of ligand in the first place. Analysis of the cumulative 59 µs of simulated data (29 µs from long trajectories and another 30 µs from 300 short trajectories, see Materials and Methods) using the MSM approach yielded estimates of the thermodynamic and kinetic parameters for the binding process which are in reasonable agreement with experiment (Table 1). The calculated kinetic on and off rate constants (*k_on_* and *k_off_*,) obtained by MSM-derived mean first passage time (*MFPT*) values (see caption of Table 1 for equations), are respectively 21 ± 9 × 10^6^ M^−1^ s^−1^ (*MFPT_on_* = 5163 ns) and 311 ± 130 s^−1^ (*MFPT_off_* = 3.2 × 10^6^ ns). As compared in Table 1, the MD-simulated *k_on_* is larger than that measured experimentally, 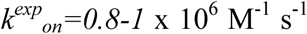. This may derive, to some extent, from the fact that the diffusion constant of the TIP3P-modelled [27] water used in the MD simulations is two to three times higher than the experimentally measured diffusion constant of water. The ligand unbinding rate constant, i.e. the so-called off-rate constant *k_off_*, computed using both MSM and Metadynamics approaches (detailed later) is in reasonable agreement with experiment 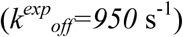. It should be emphasized that the MD estimate of *k_off_* is less precise than for *k_on_* because spontaneous unbinding events were not observed. Finally, we have also computed the standard binding free energy 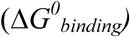 of benzene to T4L L99A using the MSM-derived stationary population of the unbound and bound conformations (see Table 2). The computed standard binding free energy of −6.9 ± 0.8 kcal/mol predicts slightly higher binding affinity than the experimentally measured 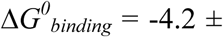 kcal/mol.

**Table 1:**
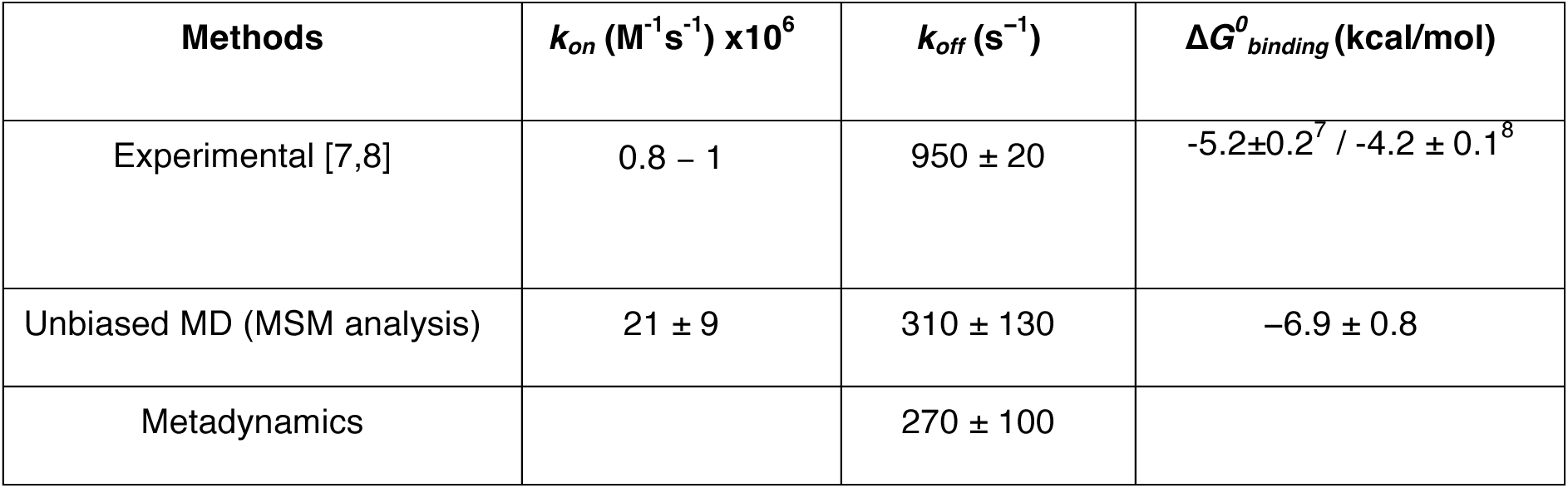
Thermodynamic and kinetic parameters estimated from MD simulations are in reasonable agreement with experimentally measured values. The thermodynamic and kinetic parameters were calculated from the four state MSM based on the mean first passage time (MFPT), *k_on_* = 1*/*(*MFPT_on_C*), *k_off_* = 1*/MFPT_off_* where C is the benzene concentration, 9.55 mM. The standard binding free energies were calculated based on the stationary populations (shown in Table 2) of bound and unbound macrostates as obtained from the MSM [20]. Unless specified, the experimental values refer to those from the NMR study by Dahlquist and coworkers [8] and are used for all comparisons.

| Methods | $k_{on} (M^{-1}s^{-1}) \times 10^6$ | $k_{off} (s^{-1})$ | $\Delta G^0_{binding} (kcal/mol)$ |
| --- | --- | --- | --- |
| Experimental [7,8] | 0.8 – 1 | $950 \pm 20$ | $-5.2 \pm 0.2^7 / -4.2 \pm 0.1^8$ |
| Unbiased MD (MSM analysis) | $21 \pm 9$ | $310 \pm 130$ | $-6.9 \pm 0.8$ |
| Metadynamics | | $270 \pm 100$ | |

**Table 2:** Properties of the four different MSM macrostates, wth MS0 and MS3 corresponding to the unbound and bound states, respectively.

| State | Population % | Lifetime (ns) | Committer Probability |
| --- | --- | --- | --- |
| MS0 | 0.11±0.06 | 2.6±4.4<br>x10<sup>3</sup> | 0.0 |
| MS1 | 6.4±3.0 x 10<sup>-3</sup> | 69±8 | 0.8±0.1 |
| MS2 | 4.7±4.5 x 10<sup>-2</sup> | 742±289 | 0.2±0.1 |
| MS3 | 99.83±0.05 | 9±4 x10<sup>5</sup> | 1 |

### Simulation trajectories reveal multiple binding pathways

One of the key findings of the current work is that there are multiple distinct pathways by which benzene binds the solvent-inaccessible cavity in T4L L99A. Contrary to the popular belief of a single dominant pathway in protein-ligand binding events,as exemplified by Shaw and coworkers in studies of GPCRs and kinases [18,19], our post simulation analyses of the six successful binding trajectories specifically identified three distinct pathways, each involving opening of channels between pairs of helices near the C-terminus of the protein. Fig 3A and supporting movies S1-S3 illustrate the three representative binding pathways via the time evolution of a benzene molecule as it reaches the binding pocket. The three pathways involve crucial conformational fluctuations of the protein and specifically, creation of crevices across certain inter-helical interfaces in the C-terminal domain. Notably, these helices were different in each of the three identified pathways as illustrated in Fig 3. For example, in trajectory 1, helices 4 and 6 temporarily move away from one another, creating a pathway to the binding site as seen in the snapshot at 7.388 *µ*s where A99 colored in orange is visible from the surface (Fig. 3B). This is in contrast to the snapshot at 7.375 *µ*s where the distance between the two helices is considerably shorter, prohibiting the entry of benzene. In trajectory 2 (Fig. 3C) benzene goes through a path created between helices 7 and 9. Benzene first associates with the surface as seen in the snapshot at 2.089 µs. The subsequent transient displacement between helices 7 and 9 creates a tunnel through which benzene enters, as shown in the snapshot at 2.090 µs. After residing at an alternate site for ~2 ns the ligand reaches the final binding site (Fig. 3C). In trajectory 3, benzene enters through an opening created at the junction between helices 5, 6, 7 and 8 (compare snapshots at 3.874 and 3.876 *µ*s, Fig. 3D). After a period of approximately 10 ns the ligand localizes to the correct binding site, as illustrated in Fig 3D.

**Fig 3:**
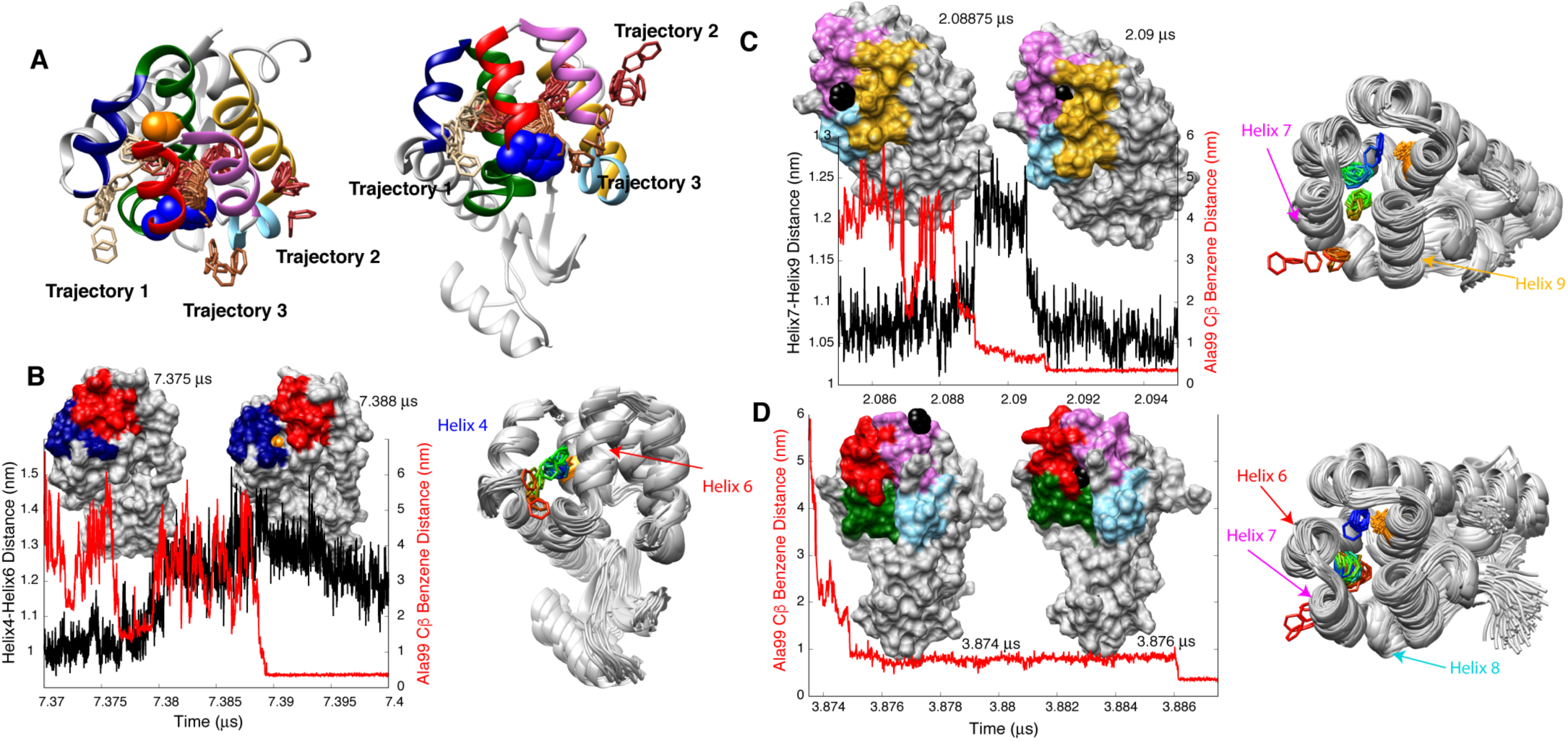
Illustration of the different pathways by which benzene reaches the binding pocket. A) Paths from three highlighted trajectories are illustrated by superimposing the backbone atoms of the protein from snapshots during the trajectories onto the crystal structure of the protein. In trajectory 1 benzene enters between helices 4 and 6, between helices 7 and 9 in trajectory 2 and through the junction region of helices 5, 6, 7 and 8 in trajectory 3. For clarity only the crystal structure of the protein is shown in A (other structures derived from the MD simulations), with Ala99 and Phe114 in orange and blue CPK representations, respectively. All helices are colored as in Fig 1. B) In trajectory 1 helices 4 and 6 transiently move away from one another, temporarily creating a pathway resulting in A99 (colored in orange) becoming visible from the surface in the snapshot at 7.388 µs. The separation between these helices returns to the starting value of ~1 nm later in the trajectory (see Fig S3). C) In trajectory 2 benzene (shown in black) enters through a pathway created between helices 7 and 9 (2.09 µs snapshot) after associating with the protein surface (2.08875 µs snapshot). D) In trajectory 3 benzene enters through an opening created at the helix 5, 6, 7 and 8 junction region. On the right side of panels B, C, D benzene entry is illustrated as in A, except that the protein is shown as a grey ribbon in each snapshot and the color of benzene changes from red to green to blue as time progresses and benzene reaches the binding site. Ala 99 is shown in orange in the structures.

In combination, these results show that key conformational fluctuations of T4L L99A lead to the formation of transient conformers where the distance between sets of helices becomes sufficiently large for benzene to reach the solvent-inaccessible cavity. As depicted in Fig S2, the spatial density profile of benzene around the protein, obtained by combining the long trajectories, provides a cumulative picture of the different pathways that lead to binding. The localization of benzene near the cavity and near the C-terminal domain helices is quite evident from the time-averaged density profile. However, we also observe certain locations near the N terminus where the ligand resides for significant periods of time (Fig S2).

Whether these highly visited locations are ultimately important for ligand binding is not clear.

### Markov state model identifies and quantifies transitions between key ligand-binding intermediates and pathways

We have used MSM-based analysis of the MD trajectories to obtain mechanistic insights into the ligand binding process. Prior to building a meaningful Markov state model, an additional three hundred, 100 ns trajectories were obtained by initiating independent simulations from different intermediates in the long trajectories. Using an aggregate set of trajectories of total duration ~59 µs (6 long trajectories + 300 short trajectories), a simple four state MSM was constructed (Fig. 4), as described in Materials and Methods. The four macrostates are illustrated in Fig 4, where MS0 and MS3 are identified with the unbound solvated benzene state and the final bound conformation, respectively, and MS1 and MS2 are two intermediates, with benzene localized near different entry points. Populations and lifetimes of each of the states, along with the committor values that measure the progress of the binding process, are summarized in Table 2. As shown in Table 2, the stationary population of the bound state, MS3, is highest and as described above, the binding free energy of −6.9 ± 0.8 kcal/mol, derived from the stationary state populations, is in reasonable agreement with the experimental measurement. The location of the benzene ligand is distinct in the intermediate structures, MS1 and MS2, highlighting again the different pathways of entry (see above). Notably, benzene is bound to different positions of the protein in the MS1 state, potentially interconverting rapidly between them. These include locations near a pair of entry points that are comprised of helices 7 and 9 or the junction between helices 5, 6, 7 and 8 that place the ligand close to the cavity. The positions of benzene in MS1 are, thus, as found in MD trajectories 2 and 3 (Figs. 3C,D). In contrast, benzene is further from the cavity in MS2, positioned instead on the surface of helices 4 and 6. MS2 is thus an intermediate of trajectory 1 that is elucidated from MD simulations (Fig. 3B). The computed committor probabilities from transition-path-theory based analysis suggest a higher (0.8) commitment of MS1 towards the bound state than that of MS2 (0.2) (Table 2). The conversion from unbound (MS0) to bound (MS3) macrostates proceeds via multiple pathways involving states MS1 and MS2. Transition path theory was used to identify paths connecting unbound and bound states and to calculate the flux through the different pathways. The kinetic pathways are illustrated in Fig 4. The contributions of the four binding pathways [MS0 → MS1 → MS3], [MS0 → MS3], [MS0 → MS2 → MS1 → MS3] and [MS0 → MS2 → MS3] to the total flux from MS0 to MS3 are 52%, 39%, 5% and 4% respectively, showing that there is no single dominant pathway. Note that the [MS0 → MS2 → MS3] path can be identified with trajectory 1, while the [MS0 → MS1 → MS3] pathway is a composite of trajectories 2 and 3.

**Fig 4:**
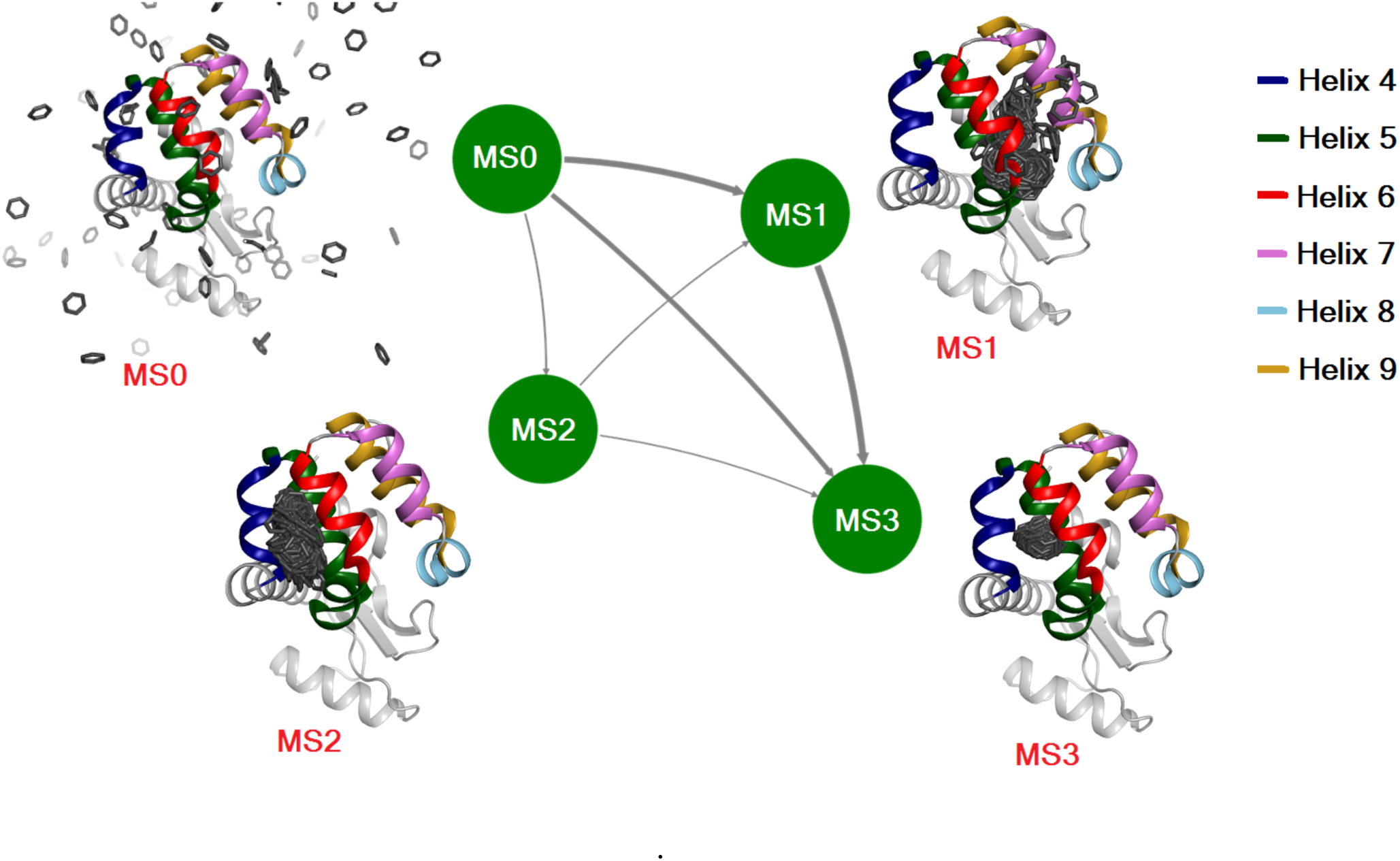
MSM analysis shows that the binding process proceeds via distinct intermediates. The MD binding data can be described by a four state (MS0-MS3) MSM network representation showing how binding proceeds via different paths through distinct intermediate states, from the free form (MS0) to the bound form (MS3). Each node represents a macrostate, with the thickness of the arrows denoting the net flux between states. 500 snapshots for each of the states are superimposed on the crystal structure of the holo enzyme (PDB 3DMX) [4], where only benzene molecules from the 500 snapshots are rendered for clarity. The population, lifetime and committor value of each of the states is shown in Table 2. Coloring is as in Fig 1 and the color-code of the helices is provided in the panel.

### Insight into the low activation barrier for ligand binding to T4L L99A

The fact that benzene is able to rapidly bind to T4L L99A is consistent with a low activation barrier. The activation free energy ∆*G*^∗^ is not readily available from experimental data, as the temperature dependencies of measured rate constants provide only an estimate of the activation enthalpy *∆H*^∗^ [28]. By assuming a diffusion controlled ligand-protein binding rate of ~10^9^ s^−1^ M^−1^, and comparing this value with the experimentally measured *k_on_* ~ 10^6^ s^−1^ M^−1^ rate, Feher et. al. estimated the free energy barrier for binding to be relatively small, 4–5 kcal/mol (6.5 to 8.5 RT) [8]. Consequently, the barrier for the unbinding process,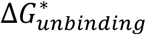, can be calculated to be ~8–9 kcal/mol (13–15 RT), based on 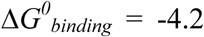 kcal/mol (Table 1). The MD simulations are also consistent with a small barrier as the observed binding rates are faster than the experimentally measured ones, even after accounting for the faster diffusion constant of the TIP3P-modeled water used here. The unbinding activation barrier can be estimated from the MD simulations using the relation 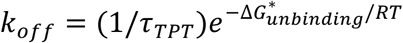 [29], where the transition path time *τ_tpt_* is the time required for benzene to transition from the cavity to the unbound form. Note that *τ_tpt_* is obtained from simulation while *k_off_ can* be measured experimentally or estimated by simulation. Values of *τ_tpt_* have been quantified directly from the six binding trajectories. In this approach, a benzene molecule was considered to be bound to T4L L99A if any of its carbon atoms was less than 0.38 nm from the Ala 99 Cβ position and, conversely, unbound when every carbon atom was greater than 0.6 nm from the Ala 99 Cβ. Based on this criterion, τ*_tpt_* values for the six trajectories vary from 0.2 to 24 ns with an average of 7 ns. Using this average value along with an experimental value for *k_off_* of 0.95×10^3^ s^−1^ [8], 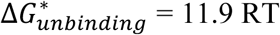 was estimated, in agreement with the barrier obtained from the analysis of Feher et al. The small value of 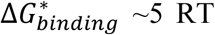, that is estimated from 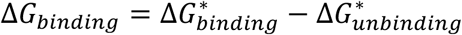, further emphasizes the small activation barrier to binding. In the above analysis barriers were estimated assuming a single binding pathway but the conclusion regarding small barriers will hold for the multiple paths as each of them occurred spontaneously during the simulations.

The MD trajectories described above provide insights into the origin of the small activation barrier for ligand binding. In all three pathways identified in this study (Fig. 3) the ligand reaches the cavity through the transient formation of pathways that are linked to the displacement of specific helices by 2–3 Å, without large scale changes in secondary structure or unfolding. The fact that T4L L99A, but not the wild-type protein, binds benzene suggests that the L99A mutation increases the probability of the protein adopting conformations that allow ligand binding. We hypothesized that the L99A mutation might increase the flexibility of the cavity-containing C terminal domain in general, leading to an increased probability for the adoption of conformations that allow ligand binding. To test this hypothesis we calculated backbone order parameters squared, S^2^, from chemical shifts for both T4L L99A and T4L WT (reerred to as 4 L WT* in what ollows, see Material and Methods) [30]. S^2^ values report on the amplitudes of ps-ns motions of amide bond vectors [31], range between 0 (isotropic motion) – 1 (rigid), and are routinely used to identify flexible regions of proteins [32,33]. The similar S^2^ vs residue profiles for both T4L L99A and WT* (Fig. 5A) indicate that fast timescale backbone dynamics are not affected by the L99A mutation, while the high S^2^ values (0.8 to 0.9) show that both forms of T4L are rigid and that rapid backbone motions are not linked directly to ligand binding. In order to ascertain whether the L99A mutation increases the population of transiently formed high energy conformers that would allow benzene to move past helices and enter the cavity we have computed free energy profiles for both T4L L99A and WT* as a function of the distances between helices 4 and 6 (Fig. 5B) and between helices 7 and 9 (Fig. 5C). Interestingly, we observe that the free energy penalty for larger inter-helical distances in T4L WT* is greater than for T4L L99A, and thus the conformations that facilitate binding are less probable for T4L WT*. For example, the probability of a Helix 4/6 separation of 1.30 nm, that is required for benzene entry (Fig. 2B, C) increases by ∼40 fold in the L99A mutant, as it is 2.21 kcal/mol more unfavorable to separate this helix pair by 1.30 nm for the WT* sequence (e^∆G/RT^ = 39 for ∆G = 2.21 kcal/mol). It is thus unlikely that a molecule of benzene could ‘squeeze’ between the helices in T4L WT* before reaching the occluded binding site due to the steric effect of residue L99. As the helices forming the C terminal domain of T4L are packed against one another, creating an opening between helices that are required for binding (Fig. 3) results in subtle positional changes to the other helices as well. A closer inspection of the trajectories showed that movement of helices 4 and 6 or 7 and 9 that is required to accommodate binding reduces the distance between side-chains of Ile78 and Ala99. A similar change in these inter-helical distances in the WT* protein would lead to a steric clash between Ile78 and the larger Leu99, as illustrated in Fig 5D in the case of Trajectory 1 where the helix 4-helix 6 distance transiently increases for only a few nanoseconds. We would like to emphasize that the population of the conformer where helices 4 and 6 are separated by 1.30 nm is extremely low, even for T4L L99A, (e^-∆G/RT^ ∼0.0046% with ∆G =3.23 kcal/mol, Fig. 5B) relative to the equilibrium conformer corresponding to a helix4-helix6 distance of 1 nm. Thus, this ligand-accessible state cannot be quantified through experiment, further emphasizing the importance of large scale MD simulations like those performed here. A similar scenario is also found for the separation of helices 7 and 9 (Trajectory 2, Fig. 3) where increasing the inter-helical distance from 1.05 nm to 1.25 nm, that is required for ligand entry, is more unfavorable by 5.9 kcal/mol for WT* than for T4L L99A.

**Fig 5:**
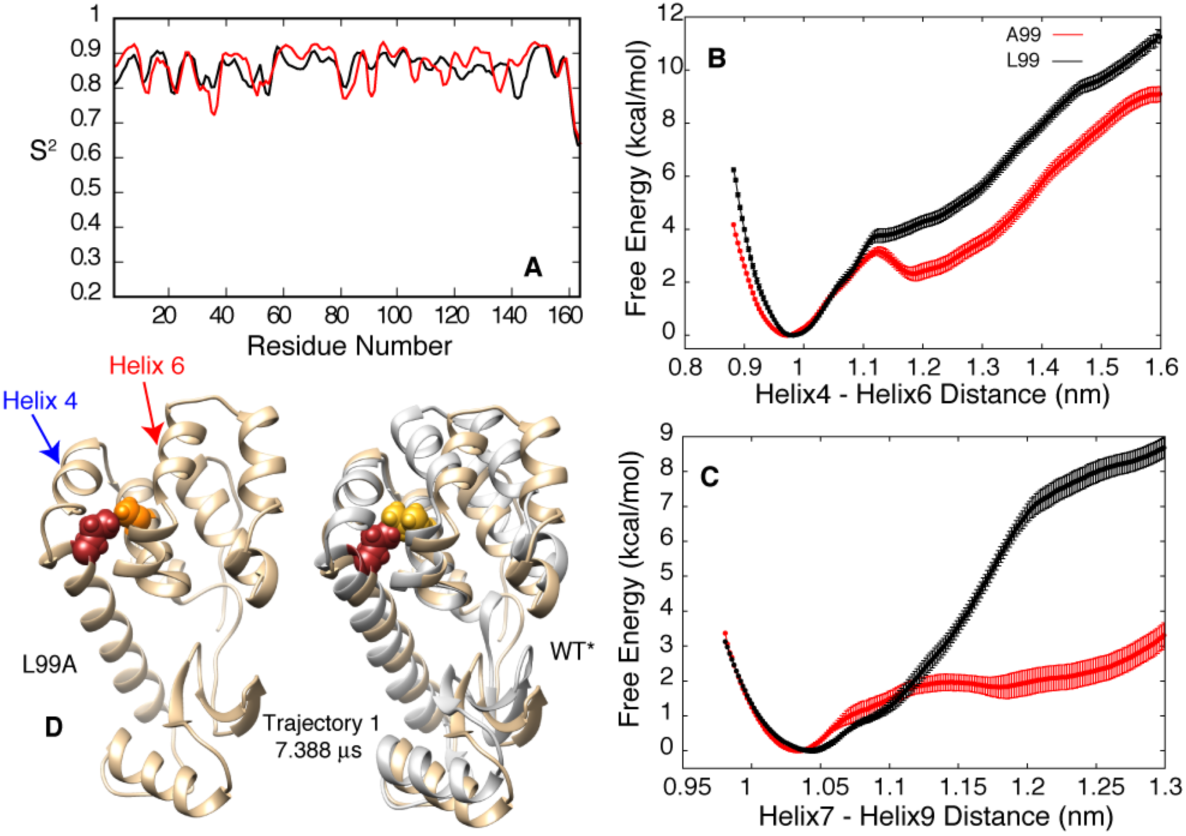
The L99A mutation stabilizes rare conformers that allow benzene to reach the cavity without substantial changes to ps-ns timescale backbone dynamics of the C terminal domain. A) NMR chemical shift derived S^2^ values [30] are very similar for T4L WT* and T4L L99A showing that the L99A mutation does not change the overall flexibility of the protein backbone on the ps-ns timescale. B,C) Free energy profiles as a function of distance between helices 4 and 6 and helices 7 and 9 showing that a larger separation between helices is more likely in the T4L L99A mutant. D) Steric clashes between Leu99 and Ile78 destabilize conformers of the WT* protein when the inter-helical distance has increased to allow benzene access to the cavity. The snapshot from trajectory 1 at 7.388 µs (L99A) has a helix 4 – helix 6 distance of 1.32 nm, with no steric clash between Ile78 (dark red) and Ala99 (orange). In contrast, in the WT* protein the larger Leu residue at the 99 position (yellow) leads to a steric clash between Ile78 and Leu99, destabilizing the conformation. The A99 to L99 mutation was performed by superimposing the backbone coordinates of residues 95 to 103 from the WT* crystal structure in grey on the snapshot from the simulation in brown.

### Unbinding trajectories validate and add to the ensemble of pathways generated from unbiased MD sampling

The computational expense of unbiased binding simulations prohibits the exhaustive survey of all of the possible kinetic pathways. Hence we explored a complementary approach of validating the identified pathways and of enriching the ensemble of binding trajectories using a recent implementation of metadynamics simulation [25] to enable the extraction of the kinetics of benzene unbinding from the cavity. Using the radial distance between the binding pocket and the ligand as the collective variable, this simulation technique infrequently deposits repulsive, history-dependent Gaussian bias along the pocket-ligand distance so as to efficiently accelerate the unbinding process. The corresponding acceleration factor, multiplied by the simulation time, provides an estimate of the time for unbinding. As depicted in Fig S4, the p-value analysis [26], as suggested by Tiwary and Parrinello, provided a good Poisson fit with a high p-value of 0.73, when averaged over numerous independent trajectories. The computed off rate, 369 s^−1^, is in close agreement with the experimentally measured off-rate, Table 1, showing that the ligand-pocket distance is a relevant collective variable for exploring benzene-T4L L99A unbinding pathways. Significantly, as depicted in Fig S3, the unbinding trajectories revealed two unbinding pathways of the ligand from the pocket. These involved benzene exiting through small openings between helices 7 and 9 and the juncture of helices 5, 6 7 and 8, pathways that are the reverse of the subset of those previously identified for binding.

## Discussions

Although ligand binding to receptors is critically important for many biological processes an atomistic understanding of how this might occur in the context of occluded binding sites is lacking. This reflects the fact that while the main biophysical tools that are used to study molecular interactions are powerful for characterizing and providing atomic resolution structures of the end points of the binding process (free and bound states) they are often much less robust in generating a description of the binding mechanism. Improvements in both computer technology and MD simulation methodology have made it possible to obtain µs-ms MD trajectories of proteins in explicit solvent so that it is possible to study processes such as ligand binding, conformational exchange, and protein folding using MD simulations [12,18,19,34]. Here, using unbiased MD simulations, we have addressed the longstanding question of how hydrophobic molecules rapidly reach buried cavities in proteins by working with a model system in which benzene binds to a 150 Å^3^ cavity mutant in T4L L99A. Central to every MD-based approach is that the force field used should be able to model the underlying free energy surface accurately. We chose to use the CHARMM force field [35] as previous MD studies had shown that the Phe114 buried to exposed conformational transition in T4L L99A and in the related protein, T4L L99A/G113A/R119P, is well modeled with it [12,36]. The structures of the benzene bound conformers obtained from the unbiased simulations are in excellent agreement with those determined by crystallography (backbone RMSD < 2 Å, Fig. 2C), with the position of the benzene within 2 Å of the binding site residues established experimentally [4]. Further, kinetic and thermodynamic parameters obtained from the simulations are in reasonable agreement with the experimental values (Table 1), providing confidence in the results of the MD study.

Our unbiased µs timescale MD simulations establish that benzene, initially fully hydrated, reaches the internal cavity created by the L99A mutation via at least three different trajectories. Notably, the process does not involve a large-scale deformation to the T4L L99A structure, nor significant changes to secondary structure, but rather small perturbations to the positions of two or more helices in the C terminal domain of the protein that create pathways for binding (Fig. 3). Similar tunnels from the protein surface to the cavity were also observed in a 30 µs MD simulation of T4L L99A using the AMBER force field [37]. The existence of multiple pathways coupled with a small net activation barrier explains how benzene can bind the occluded cavity rapidly.

Previously we had used combined MD simulations and NMR relaxation experiments to study how the side-chain of residue Phe114 fills the cavity created by the L99A mutation, that in some sense serves as a surrogate to hydrophobic ligands such as the benzene molecule considered here. It might be expected, therefore, that the binding pathways of Phe114 and benzene would share some similar features. Notably, in one of the trajectories (Trajectory 3) benzene enters the cavity in a manner similar to Phe114 [12] from its solvent exposed conformation.

The binding of ligands to occluded sites in other proteins, such as oxygen binding to the buried heme group in myoglobin, has been studied using a variety of experimental and computational techniques [38]. In the case of myoglobin, the mechanism of binding also involves small barriers, multiple pathways and secondary binding sites in the protein [38]. The plasticity of T4L L99A allows the larger benzene molecule to reach the cavity in a manner similar to how oxygen connects with the heme group in myoglobin. The origin of the conformational plasticity in T4L L99A that facilitates this process is, in fact, the L99A mutation itself (Fig. 5). Replacing the Leu99 residue with the smaller Ala not only creates the cavity but also allows helices to move with respect to one another to create the pathways that are required for binding. In contrast, the presence of Leu99 destabilizes the binding competent conformations by clashing with Ile78 (Fig. 5). It remains of interest to investigate if the benzene binding rate can be modulated by mutating Ile78 to smaller hydrophobic residues. Similar to the binding of benzene, protein plasticity and cavities have also been implicated in oxygen binding to T4L L99A [14].

Preliminary observations into the mechanism of ligand unbinding from T4L L99A have recently been published and are in agreement with the results of this study. Kitahara et. al. [14] developed a simple and elegant ^15^ N NMR based method to detect O_2_ binding sites in proteins using T4L L99A as a model system. In order to quantify the ^15^ N chemical shift changes that were observed upon oxygen binding MD simulations were performed to understand the rotational and translational diffusion properties of oxygen molecules in the T4L L99A cavities. Notably, an oxygen molecule escaped from the cavity via a pathway that is the reverse of Trajectory 1 (Fig. 3). In MD simulations performed to understand water filling the cavity at high pressures, water molecules in the cavity escaped from the cavity due to transient openings formed at the junction of helices 5, 6 and 7 [39] and the same helices transiently move to create a pathway to the cavity here in Trajectory 3. Wang et. al. used biased simulations to map out the free energy surface of T4L L99A along the pathway by which Phe114 transitions into the L99A cavity. In some of their adiabatic-biased MD simulations of benzene unbinding a pathway that is the reverse of Trajectory 3 was observed [36]. In a separate 27 µs MD trajectory of T4L L99A performed to understand the Phe114 exposed to buried conformational transition, tunnels leading to the cavity from solvent were transiently formed [37], as observed in the present work. A recent implementation of a biased simulation approach by Lindorf-Larson and coworkers also has extracted the kinetics of the benzene recognition process [40]. McCammon and coworkers[41] have employed biased simulation approach to explore thermodynamics of benzene binding to cavity. However, none of these simulation-based studies address the mechanism by which ligands *spontaneously* bind to buried protein cavities and consequently insights into multiple pathways and low free energy barriers were not obtained.

Recent studies suggest that diverse processes such as protein folding and protein interconversion between different compact conformers proceed via multiple pathways and/or small barriers [12,42,43]. More studies are required to determine if small activation barriers and multiple paths are common features of biomolecular recognition processes and to characterize at atomic resolution what these pathways are and the possible role of high pressure on cavity solvation[39].

## Materials and Methods

Here, as in most biophysical studies involving T4Lysozyme, the cysteine free version of the protein [44] was used, T4L WT*, where Cys54 and Cys97 are replaced by Thr and Ala, respectively. In T4L L99A the L99A mutation is introduced into the T4L WT* background.

## MD simulations and analysis

### Unbiased binding simulations

The X-ray crystallographic structures of T4L L99A free and benzene bound forms (PDB code: 3DMX and 3DMV) were used as the starting points for all of the simulations [4]. All binding simulations were initiated by placing the apo form of L99A T4L at the center of a cubic box of dimension 7.2 nm with the empty space then filled with water and ion molecules. The system was solvated with 11613 triatomic water molecules and sufficient numbers of sodium and chloride ions were added to keep the sodium chloride concentration at 150 mM and render the system charge neutral. Two benzene molecules, corresponding to a concentration of 9.55 mM, that is less than the benzene solubility limit in water (20 mM) [45], were placed in random positions in the solvent. The two molecules were allowed to diffuse freely and no artificial bias was introduced throughout the simulation. The system included a total of 37541 atoms. The protein, benzene and ions were parameterized with the charmm36 force field [35] and the TIP3P water model [27] was used.

All MD simulations were performed with the Gromacs 5.0.6 simulation package [46], in most cases benefiting from usage of Graphics processing units [47]. During the simulation, the average temperature was maintained at room temperature (303 K) using the Nose-Hoover thermostat [48,49] with a relaxation time of 1.0 ps and an average isotropic pressure of 1 bar was maintained with the Parrinello-Rahman barostat [50]. The Verlet cutoff scheme [51] was employed throughout the simulation with the Lenard Jones interaction extending to 1.2 nm and long-range electrostatic interactions treated by Particle Mesh Ewald (PME) summation [52]. All bond lengths involving hydrogen atoms of the protein and the ligand benzene were constrained using the LINCS algorithm [53] and water hydrogen bonds were fixed using the SETTLE approach [54]. Simulations were performed using the leapfrog integrator with a time step of 2 fs and initiated by randomly assigning the velocities of all particles.

Initially six long independent unbiased trajectories were performed, differing in the assignment of initial velocity seeds. These ranged between 2.5 and 8.1 µs with a total simulation length of 29 µs. The simulations were only terminated after one of the two copies of the ligand settled into the target binding site. The binding process was ascertained via inspection and by checking (i) the radial distance between respective centers of mass of benzene and the cavity and (ii) the root-mean-squared deviation (RMSD) of the simulated conformation of the protein-ligand system from that of X-ray crystal structure of holo T4L L99A (PDB id: 3DMX) [4] that included benzene atoms and those of the protein cavity. The ‘cavity’ or the ‘binding pocket’ was defined by the protein heavy atoms within 5Å of benzene in the X-ray structure of holo T4L L99A (PDB id: 3DMX). For the calculation of RMSD, the cavity plus ligand of each frame of the trajectory was first translationally and rotationally aligned with that of the crystal structure of the benzene-bound form of protein (PDB id: 3DMX) and then the deviation was computed. The ligand was ascertained to be bound to the cavity when the RMSD remained below 0.5 nm and the cavity-ligand distance was below 0.2 nm for a simulation duration of at least 100 ns.

Apart from the six µs-long trajectories discussed above, we have also performed 300 short independent trajectories, each 100 ns in duration. These short trajectories were initiated from different intermediates observed from the long binding simulations and improved the statistics of the MSM that was derived from analysis of the MD trajectories (see below). Overall, an aggregate of about 59 µs of unbiased trajectory was recorded.

### MSM analysis of the MD data

The generation of a MSM is a powerful way to automatically identify the kinetically relevant states and their interconversion rates from the simulated trajectories [55–58]. We employed PyEMMA [59] (http://pyemma.org) to construct and analyze the MSM obtained from the combination of all the recorded MD trajectories. The nearest-neighbor (cut-off 0.5 nm) heavy-atom contacts between benzene and protein residues were used as input coordinates. The time-lagged or time-structure-based independent component analysis (tICA) [60–62] with a correlation lag time of 20 ns was used for dimensionality reduction, which linearly transforms the high dimensional input data into a slow linear subspace (i.e. collective coordinates sorted by ‘slowness’). The high dimensional input coordinates were subsequently projected onto 12 tICA components, covering more than 95% of the kinetic variance. We employed the k-mean clustering algorithm [63] to discretize the tICA data into 100 clusters. Then 100 microstates MSMs were constructed at variable lag times to identify the appropriate lag time. A lag time of 10 ns was used to construct the final MSM as the implied time scale leveled off at about 10 ns which ensures the Markovianity of the model. To better understand the dynamics of ligand binding a coarsegrained kinetic model with four metastable states was constructed using a hidden Markov model (HMM) [64].

### Free energies of helix opening

The umbrella sampling simulation technique [65] was used to map the free energy surface underlying the opening of a pair of helices to facilitate ligand binding to the cavity in T4L L99A. To this end, we performed two independent umbrella sampling simulations where the distance between centers of mass of Cα-atoms of helices 4 and 6 or of helices 7 and 9 was used as the collective variable (CV). The value of the CV ranged from 0.9 nm – 1.6 nm for the distance between helices 4 and 6 and from 1 to 1.3 nm for helices 7 and 9. The starting configuration for each window was picked from the prior unbiased simulation trajectories, with a spacing between consecutive windows of 0.05 nm. Each window was restrained to the desired value of the CV using a harmonic potential of force constant 8000 kJ/mol/nm^2^ to ensure that the distribution of the CV for each window remains Gaussian with sufficient overlap in the distribution of two consecutive windows so that the entire space of the CV is sampled. Each of the windows was sampled by the umbrella sampling technique for 20 ns. Finally, the weighted histogram analysis method (WHAM) [66,67] was employed to reweight the umbrella-sampled windows and to obtain the unbiased potential of mean force as a function of the CV. In order to investigate the effect of the L99A mutation on the free energetics of helix opening, the entire series of umbrella samplings was repeated by in-silico mutation of Ala99 in each of the starting configurations to Leu99.

### Unbinding simulations using an enhanced sampling technique

The recent implementation of metadynamics by Tiwary and Parrinello [25] was used to compute unbinding rates of benzene from the hydrophobic cavity of L99A T4L and to extract unbinding pathways. In this novel method, by making the bias deposition slower than the time in bottlenecks, and by using a suitable CV capable of distinguishing between bound and unbound states, it is possible to keep the transition state relatively bias-free through the course of the metadynamics simulation.

This so-called ‘infrequent metadynamics’ approach allows one to accelerate the transition from ligand-bound to unbound states, without affecting the transition state. More importantly, this technique provides an estimate of the acceleration factor, which when multiplied with metadynamics simulation times, provides the true unbinding time. As elaborated in the original work of Tiwary and Parrinello [25], the acceleration factor is given by the time *α* = 〈*e^*ßv(s,t)*^*〉*_t_* where *s* is the CV being biased, *ß* is the inverse temperature, *V(s,t)* is the bias experienced at time *t* and the subscript *t* indicates that averaging is performed under the time-dependent potential. The above expression is valid even if there are multiple intermediate states and numerous alternative reaction pathways. In our current work, we employed the distance between the centers of mass of the cavity and benzene as the CV using a Gaussian width of 0.025 Å. One of the reasons for choosing the distance between the binding pocket and the ligand as the CV was that it can distinguish between bound and unbound states very well, while providing an estimate of binding free energies that agree with those obtained from experiment. During the ‘infrequent metadynamics’ simulation approach, the Gaussians were deposited every 10 ps with a starting height of 1.2 kJ/mol and gradually decreased on the basis of a well-tempered metadynamics biasing factor, Γ = 6. A total of 15 such ‘infrequent metadynamics’ simulations having different initial velocity distributions were spawned with the ligand fully bound to the cavity and simulations were stopped only when the ligand became fully unbound, reaching bulk-solvent i.e. the distance between the binding pocket and the ligand exceeded 3 nm. Acceleration factors, as described earlier and elaborated on in reference [25], were computed during the simulations and multiplied with the Metadynamics run time to get the true unbinding time.

The transition time statistics so obtained were then subject to a Poisson analysis [26] to ascertain their reliability (based on the Kolmogorov-Smirnov statistical analysis) and to obtain the average unbinding time, from which the unbinding rate is easily calculated. Our approach of performing metadynamics simulations using the cavity-ligand separation as a single CV was sufficient to pass the so-called p-value test which suggested that the frequency of bias was slow enough and the CV was appropriate to accelerate unbinding. All simulations were carried out using Gromacs 5.0.6 patched with the PLUMED plugin [68].

## NMR experiments and analysis

### NMR Samples

^15^ N,^13^ C labeled T4L WT* and T4L L99A were overexpressed and purified as described previously [69]. The protein concentration in the NMR samples (50 mM sodium phosphate, 25 mM NaCl, 2 mM EDTA, 2 mM NaN_3_, pH 5.5, 10% D_2_O buffer) was ∼1.5 mM.

### NMR experiments

Assignments for T4L WT* were obtained using standard triple resonance experiments [70,71], with assignments for T4L L99A published earlier [72]. All experiments were performed at 22°C on a 700 MHz Bruker Avance III HD spectrometer equipped with a triple resonance cold probe.

### NMR data processing and analysis

NMR data were processed using the NMRPipe software package [73] and the resulting spectra were visualized and analyzed using SPARKY [74]. To overcome complications due to chemical exchange, order parameters were estimated from chemical shifts according to the method of Berjanskii and Wishart [30], as implemented in TALOS+ [75].

## Acknowledgements

This work was supported by startup funds from Tata Institute of Fundamental Research, Hyderabad (TIFRH) (JM and PV), early career research funds provided by Department of Science and Technology of India (ECR/2016/000672) (JM) and the Extreme Science and Engineering Discovery Environment (XSEDE) [TG-MCA08X002] (JM).

## Supporting Information

**S1 File.** PDF file containing four supporting figures including Fig S1, S2, S3 and S4.

**S1 Movie.** Movie (see uploaded media file titled “S1_movie.mp4”) highlighting trajectory 1. Benzene binds to the cavity via the gateway between helices 4 and 6, created by transient distance fluctuations between these elements of structure. The protein is shown in cartoon representation, the cavity by a solid surface representation and the benzene molecule via a licorice representation.

**S1 Movie.** Movie (see uploaded media file titled “S2_movie.mp4”) of trajectory 2 showing benzene binding via a pathway between helices 7 and 9 created by transient helix 7–9 distance fluctuations. The protein is shown in cartoon representation, the designated cavity by a solid surface representation and the benzene molecule via a licorice representation.

**S3 Movie.** Movie (see uploaded media file titled “S3_movie.mp4”) of trajectory 3 whereby benzene enters T4L L99A through the junction region of helices 5–8. The protein is shown in a cartoon representation, the designated cavity by a solid surface representation and the benzene molecule via a licorice representation.

